# FtsK in motion reveals its mechanism for double-stranded DNA translocation

**DOI:** 10.1101/828319

**Authors:** Nicolas L. Jean, Trevor J. Rutherford, Jan Löwe

**Affiliations:** MRC Laboratory of Molecular Biology, Cambridge, UK

**Author notes:** Correspondence to: Jan Löwe. MRC Laboratory of Molecular Biology, Cambridge CB2 0QH, UK.

## Abstract

FtsK protein contains a fast DNA motor involved in bacterial chromosome dimer resolution. To understand how FtsK moves DNA, we solved the 3.6 Å resolution cryo-EM structure of the motor domain of FtsK while translocating double-stranded DNA. Each subunit of the hexameric ring adopts a unique conformation and one of three nucleotide states. Two DNA-binding loops within four subunits form a pair of spiral staircases within the ring, interacting with the two DNA strands. This suggests that simultaneous conformational changes in all ATPase domains at each catalytic step generate movement through a mechanism related to filament treadmilling. While the ring is only rotating around the DNA slowly, it is instead the conformational states that rotate around the ring as the DNA substrate is pushed through.

## Main Text

DNA and RNA motors are a broad family of proteins that convert chemical energy from nucleotide triphosphate hydrolysis into motion relative to nucleic acids. Among them are ring-shaped helicases and translocases, which belong to the AAA+ and RecA families in eukaryotes and prokaryotes, respectively (*1*). With a translocation rate of up to 17.5 kb.s^−1^ (*2*), the bacterial FtsK protein contains the fastest translocation activity known. FtsK is an essential part of the bacterial cell division machinery, recruiting downstream factors through its N-terminal trans-membrane domain (*3*, *4*). Its C-terminal cytoplasmic domain, which follows a long linker domain can be subdivided into 3 modules: α, β and γ, with β adopting the RecA-fold with ATPase activity. It is the C-terminal αβγ domain that is involved in chromosome segregation and dimer resolution (*5*–*7*). Upon recognition by the γ module of a direction-determining sequence in the genome, named KOPS (FtsK orienting polar sequences) (*8*–*12*), α and β oligomerize as a hexameric ring around double-stranded DNA (forming FtsK_αβ_-dsDNA) (*13*). FtsK_αβ_ then starts pumping DNA, translocating towards the chromosome terminus where the γ module activates recombination by recombinases such as XerCD (*14*, *15*). Previously, FtsK_αβ_ has been crystallized as a 6-fold symmetrical ring (*13*), but without dsDNA in the structure, the mechanism by which it translocates has remained unclear. Structures of DNA- or RNA-helicase complexes have suggested that nucleic acids, similarly to peptidic substrate in unfoldases, are generally recognized by a more-or-less asymmetrical ring through motifs organized into a partial helix, akin to a spiral staircase (*16*–*21*). However, the proposed translocation mechanisms for helicases are based on single-stranded nucleic acids, and it remains to be seen if similar mechanisms apply to motors translocating on more rigid double stranded DNA substrates.

Here, we solved by electron cryo-microscopy (cryo-EM) structures of FtsK_αβ_ from *Pseudomonas aeruginosa* bound to dsDNA. Purified FtsK_αβ_ mixed with 45 bp dsDNA displays ATPase activity that is too fast for EM grid preparation and 1D ^31^P NMR (Figs 1A and S1), as expected from its previously determined turnover rate of 2,600 ATP s^−1^ per hexamer (*22*). In contrast, FtsK_αβ_ hydrolyses ATPγS (but not AMPPNP) at a much lower rate (at least three orders of magnitude lower) (Figs. 1A and S1). Cryo-EM with DNA, in absence or presence of the nucleotides ADP, AMPPNP and ATPγS confirmed that FtsK_αβ_ forms hexameric rings without the γ module (Fig. 1B), as observed previously (*13*). Although not visible in micrographs directly due to its short length, the 45 bp dsDNA can be detected after 2D classification inside the central pore (Fig. 1B). Cryo-EM maps of the nucleotide-free, ADP-, and AMPPNP-bound FtsK_αβ_-dsDNA complexes were obtained at resolutions of 4.91 Å, 4.63 Å and 4.80 Å respectively (Fig 1C and table S1,). In these non-translocating states, the FtsK_αβ_ rings remained symmetrical, with uniform nucleotide states around the ring, and no conformational differences relative to the previous crystallographic structure being obvious (Figs. 1C and S2). Presumably, the DNA’s almost central position in the pore, in conjunction with the almost perfect αβ ring symmetry, led to poor alignments and thus some loss of map quality around the DNA since it does not have six-fold symmetry (see Supplementary Materials and Methods). In the symmetrical nucleotide-free, ADP- and AMPPNP-bound structures most contacts with the dsDNA seem to occur at the N-terminal α-ring and the base of the β subdomain where DNA enters the pore.

**Fig. 1:**
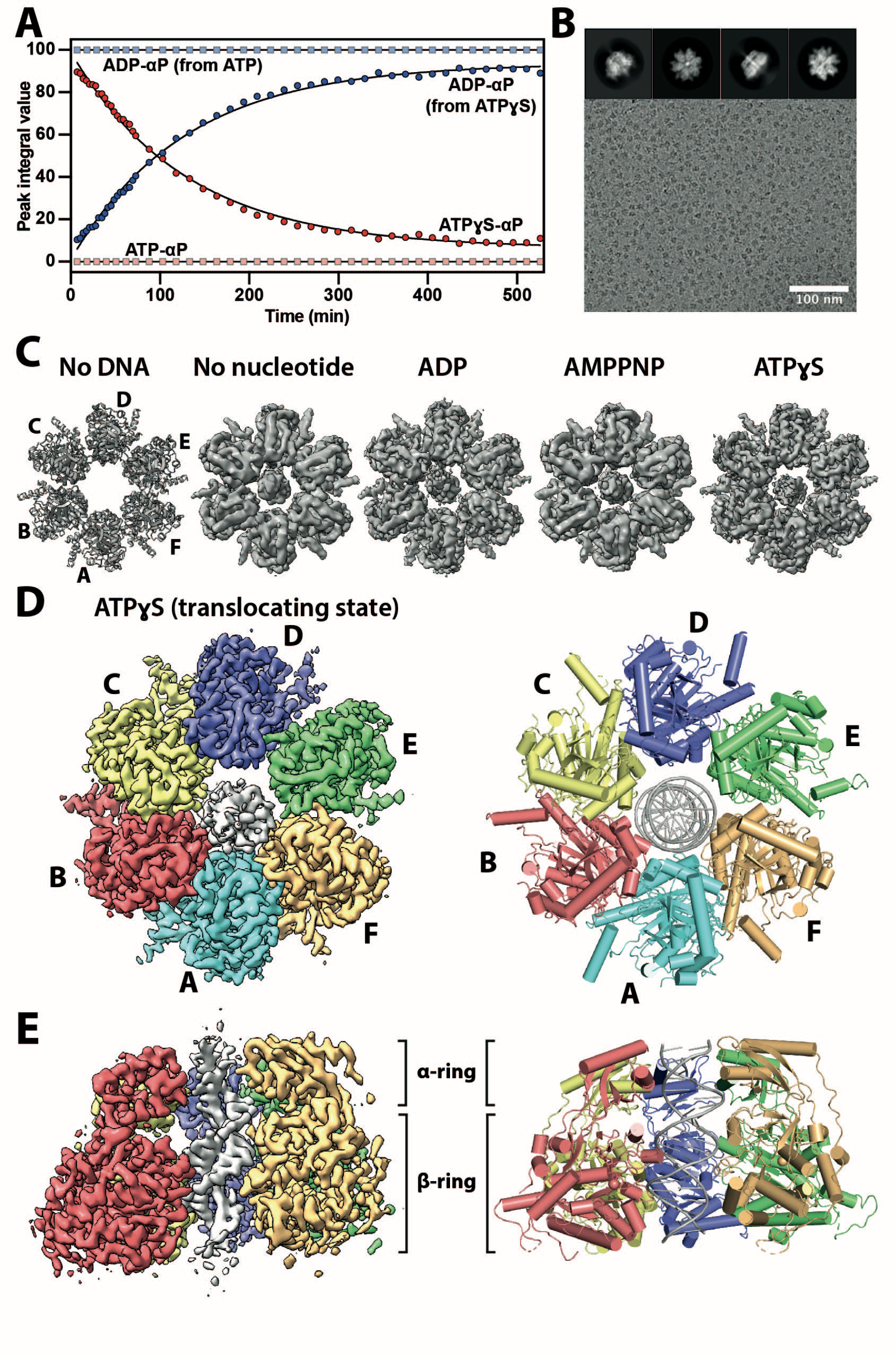
Cryo-EM structures of the FtsK_αβ_-DNA complex. (**A**) ATP and ATPγS hydrolysis by FtsK_αβ_ over time. The data points for the α-phosphates of ATP (light red) / ATPγS (dark red) / ADP (light and dark blue) were determined from their peak integrals obtained by 1D 31P NMR experiments. ATP is fully hydrolysed into ADP before the first data point, seven minutes after nucleotide addition. (**B**) Representative cryo-EM micrograph of FtsK_αβ_-dsDNA + ATPγS. Typical 2D classes (top). **(C)** Top views of FtsK_αβ_ crystal structure (PDB 2IUU) and FtsK_αβ_-dsDNA cryo-EM maps determined here. Only map III.E (translocating state, see Fig. S3) is shown for ATPγS. **(D)** and **(E)** 3.6 Å resolution cryo-EM map (left) of the FtsK_αβ_-dsDNA + ATPγS complex and refined atomic model (right). Each subunit is coloured differently, with the double-stranded DNA in the pore. Shown from the α-ring where the DNA exits the pore during translocation (D) and as side view with subunit A removed (E).

The FtsK_αβ_-dsDNA-ATPγS sample was much more heterogeneous, the particles less symmetrical, and DNA generally much better defined after 3D refinement. Overall, three main particle classes could be distinguished, II.A, III.B and III.E, accounting each, with their other equivalent classes, for 30.0%, 17.9% and 25.4% of total particles selected after 2D classification, respectively. After pruning further for highest quality and resolution, these numbers dropped to 4.9%, 4.3% and 5.2%, respectively (Fig. S3). States II.A and III.B were reconstructed to 3.99 Å and 4.34 Å resolution and are in very similar conformations (Figs. S4, S5 and Table S2). Both structures are moderately asymmetric, mostly due to a movement of the β subdomains in subunits A and B, turned away from the α-ring (Fig. S5). Other subunits are again very similar to the previous crystallographic structure. The main difference between these two states lies in the length of the dsDNA, which is 5 bp shorter in II.A on the side closest to the β subdomain, at the pore entry. From the maps we determined that in these structures ADP occupies all nucleotide-binding pockets and this suggests that they represent stalled states, with II.A having reached the end of the short 45 bp DNA after translocation.

Particles in the final class, III.E, were reconstructed into a map at 3.65 Å resolution (Fig. S6 and table S2) and result in a highly asymmetric ring with mixed nucleotide occupancy that is also slightly tilted relative to the DNA’s longitudinal axis (Figs. 1D, 1E, S4). For reasons explained below, we propose that class III.E represents the protein while it is actively translocating on the DNA, and from here on refer to it as FtsK_αβ_.

In FtsK_αβ_, solved under ATPγS turnover conditions, each of the subunits A-F adopts a different conformation, named here 1 to 6 (Fig. 2A). The orientations and positions of the β subdomains relative to the α subdomains vary around the ring, describing mostly a planar rocking motion approximated by the angle between the S383-I365-K657 Cαatoms. The subdomains are closest to each other in conformation 6, with a measured angle of 33.5°, similar to the crystallographic structure. Going clockwise around the ring, β moves away from αand also towards the adjacent subunit on its left. A maximum is reached in conformation 3 with an angle of 69.4°, after a gradual increase through conformations 1 and 2. The ATPase subdomain finally resets to conformation 6 through conformations 4 and 5. Therefore, the inter-subdomain angles describe a conformational wave (Fig 2A, bottom). The conformational diversity within the subunits of the FtsK_αβ_ hexamer results in an asymmetrical β-ring, that also creates a significant variation in inter-subunit packing, as highlighted by the 65% decrease in buried subunit interface area between interfaces 2-3 and 5-6 (Fig. S7). The subunit packing in conformations 5 and 6 might be loose enough to enable some freedom in β movement without compromising the integrity of the ring and their lower resolutions support this hypothesis. In contrast, symmetry in the α-ring is mostly preserved and it is like a rigid aperture (Fig. S7).

**Fig. 2.**
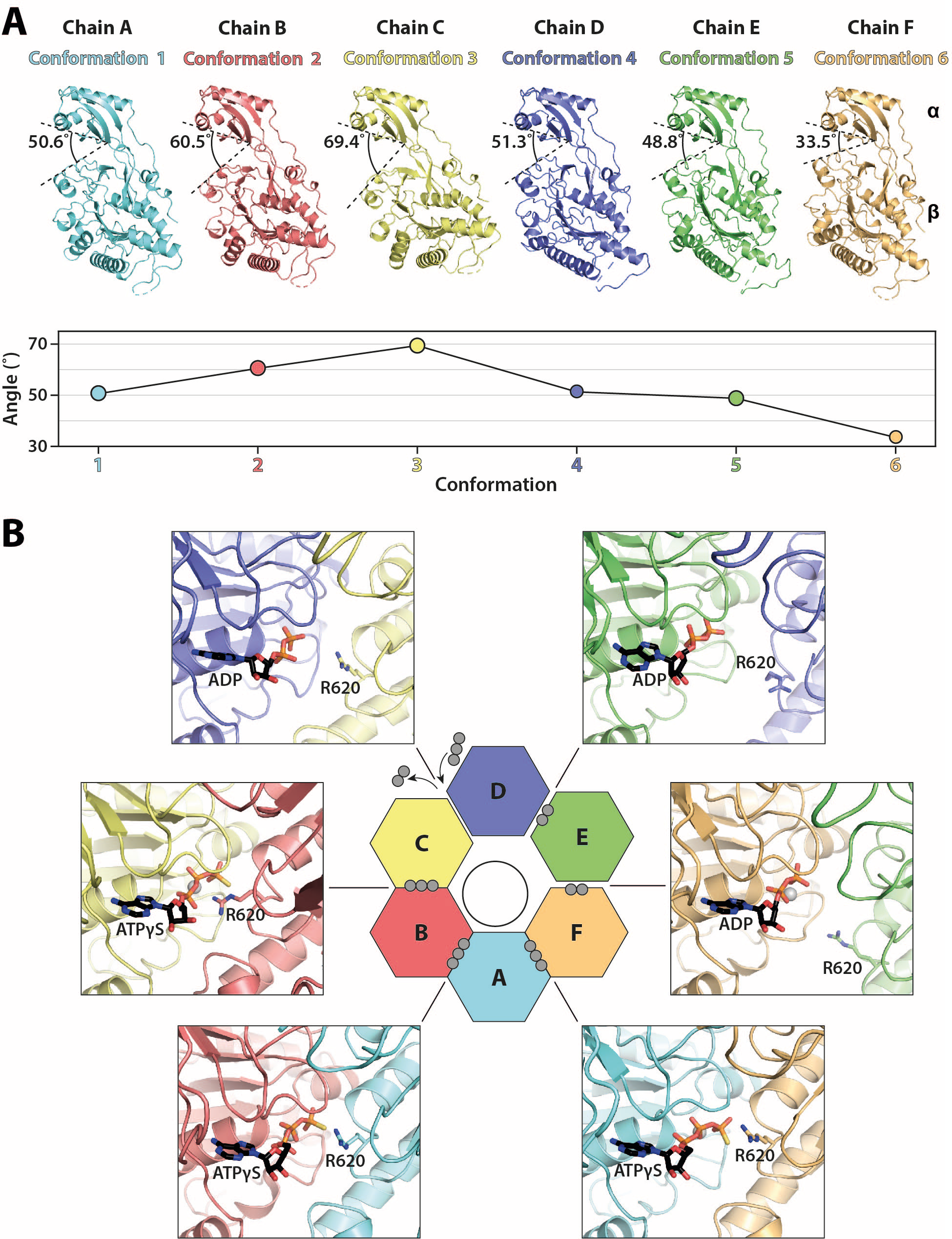
In the translocating state, FtsK_αβ_ subunits around the ring undergo a continuous conformational transition. **(A)** Side-by-side comparison of FtsK_αβ_ subunits aligned on the α-subdomains. Each subunit is in a different conformation, from here on identified by its colour. The angle between α and β subdomains (between S383-I365-K657 Cα) is indicated and plotted in the graph below to highlight the wave nature of the conformational differences. **(B)** Nucleotide states around the FtsK_αβ_ ring in ATPase sites that are located at the interfaces between two neighbouring subunits. Although ADP is modelled in the C-D pocket, inspection of the EM map suggests that it is likely in exchange with ATPγS. The activity-controlling arginine finger (R620) is highlighted. Grey spheres indicate phosphates per nucleotide.

The conformational diversity within the FtsK_αβ_ ring is correlated with three different nucleotide states around the β-ring (Figs. 2B and S8). In RecA-like hexameric translocases, active site pockets are located at the subunit interfaces. In FtsK_αβ_, ATPγS is well-resolved at the F-A, A-B, and B-C interfaces (between conformations 6-1, 1-2 and 2-3). In the ATPγS-bound state, the subunit packing allows the arginine finger (R620) from one subunit to reach the nucleotide in the pocket of its adjacent subunit. Two other interfaces, D-E and E-F (conformations 4-5 and 5-6) clearly show map densities for only two phosphates of ADP molecules, generated through in-situ ATPγS hydrolysis. In agreement with a post-hydrolysis state, here the arginine finger is located away from the nucleotide pockets. However, the two interfaces show subtle variations, such as an Mg^2+^ ion coordinated only in the E-F pocket and a much higher atomic B-factor for the ADP in the E-D pocket (90.1 Å^2^ versus 58.7 Å^2^). Finally, the nucleotide pocket at the C-D interface (conformations 3-4) shows less well-defined nucleotide density for the two ADP phosphates. An additional density for a γ-phosphate, visible only at lower thresholds, suggests that the pockets in the particles averaged are likely in exchange between ADP and fresh ATPγS. In agreement with such a transitional state, the arginine finger lies in a position mid-way between the ATPγS and ADP states.

The DNA density is very well defined for 20 base pairs, traversing the pore from one end to the other. Density for the nucleobases is the result of averaging over all four possibilities because FtsK’s DNA translocase activity is not sequence specific, and hence no sequence was assigned to the DNA. Additional map density extended on both sides but had lower quality presumably due to DNA flexibility outside the pore. Contrary to the nucleotide-free, ADP- and AMPPNP-states, in the translocating state DNA makes extensive contacts with four subunits in the β-ring, F, A, B and C, adopting conformations 6, 1, 2 and 3 (Fig. 3). The most prominent contacts with all four subunits involve basic residues protruding into the pore from two loops contacting the phosphodiester backbone on both strands of the minor groove. Loop I binds to two phosphates from one strand, exclusively through contacts facilitated by K657 and R661. Similarly, two phosphates are recognized by loop II, through R632 and to a lesser extent K643. H-bonds with additional loop II residues S634, V635 and G640 reinforce this interface. The conformational heterogeneity positions these loops in two helical paths, or ‘spiral staircases’, following the DNA’s helix precisely (Movie S1). This arrangement is reminiscent of the single spiral staircases observed in ring-shaped helicases (*23*). In FtsK, conformation 3 (subunit C) produces a DNA interface at the bottom of the staircase, followed by the two other, ATPγS-bound conformations 2 and 1 (subunits B and A). The path is terminated by the ADP-bound conformation 6 (subunit F) at the top. Conformations 5 and 4 (subunits E and D) are not bound to DNA, allowing two subunit positions for the gradual transition from the top of the spiral staircase to the bottom, resetting the subunit conformation. The gradual decrease in α to β angle from conformation 3 to 1 creates a smooth transition, which is broken somewhat by conformation 6 and its larger step size. Curiously, this irregularity distorts the double helix by widening the minor groove by up to 25% (from 11.8 Å to 14.7 Å) from canonical B-form DNA (Fig. S9A).

**Fig. 3.**
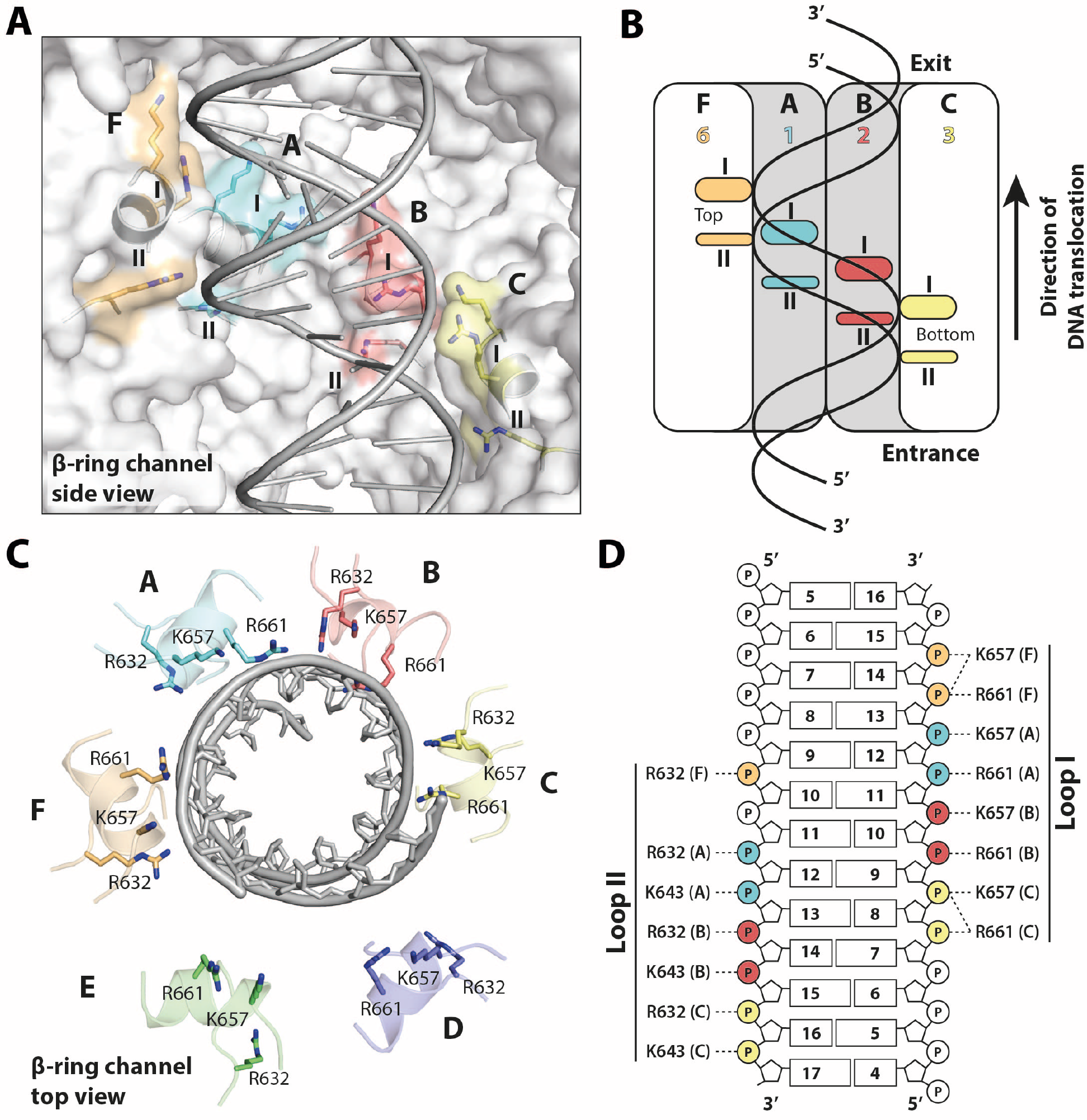
Interactions with double-stranded DNA within FtsK’s translocating pore. **(A)** Recognition by DNA-binding loops organized into a double helical arrangement, a ‘double spiral staircase’. Basic residues interacting with the phosphodiester DNA backbone are highlighted. Subunits D and E, whose β-subdomains do not interact with DNA have been removed for clarity. **(B)** Schematic representation of (A). **(C)** Top view of (A) with all six subunits. Subunits D and E are far away from the DNA. **(D)** All direct interactions between DNA and basic residues in loops (staircases) I and II. Each of the four interacting subunits contacts two phosphates, implying a translocation of exactly two base pairs per step.

Several contacts are made with the DNA at the pore’s exit, at the position of the α-ring (Fig. S9B). These interactions are mediated by interactions with K377 and R380. However, because of the largely preserved rotational symmetry of the α-ring, these residues form a flat ring, almost perpendicular to the DNA’s long axis. This configuration prevents tracking of the DNA helicity, but might allow subunits on both sides of the DNA to maintain its alignment with the pore’s axis as it is exiting. Indeed, one DNA strand is recognized by protomers in conformations 1 and 6 (subunits A and F), while the other is by conformation 4 (subunit D), whose β subdomain is disengaged from DNA.

Because the subunits seen in the asymmetric ATPγS FtsK_αβ_ structure appear to describe an entire ATP hydrolysis and conformational cycle, the structure directly suggests a mechanism for dsDNA translocation by FtsK_αβ_ that involves rotating the conformational states around the ring. Hence our mechanistic model has at its centre a concerted, simultaneous conformational change of all subunits. At each step, all subunits change conformation and advance along the nucleotide hydrolysis cycle by 1/6^th^ of the entire cycle, and more-or-less simultaneously (Fig. 4A and Movie S2). The nucleotide states around the ring clearly indicate that the reaction cycle advances around the ring clockwise (Fig. 4A), as postulated previously (*13*). Conformation 1 (subunit A in the structure) is therefore competent for ATP hydrolysis. Importantly, given the right-handedness of the DNA, this leads to translocation of the DNA from the bottom of the β-ring to the top of the α-ring (Fig. 4B) and this is in agreement with previous data (*10*, *13*).

**Fig. 4.**
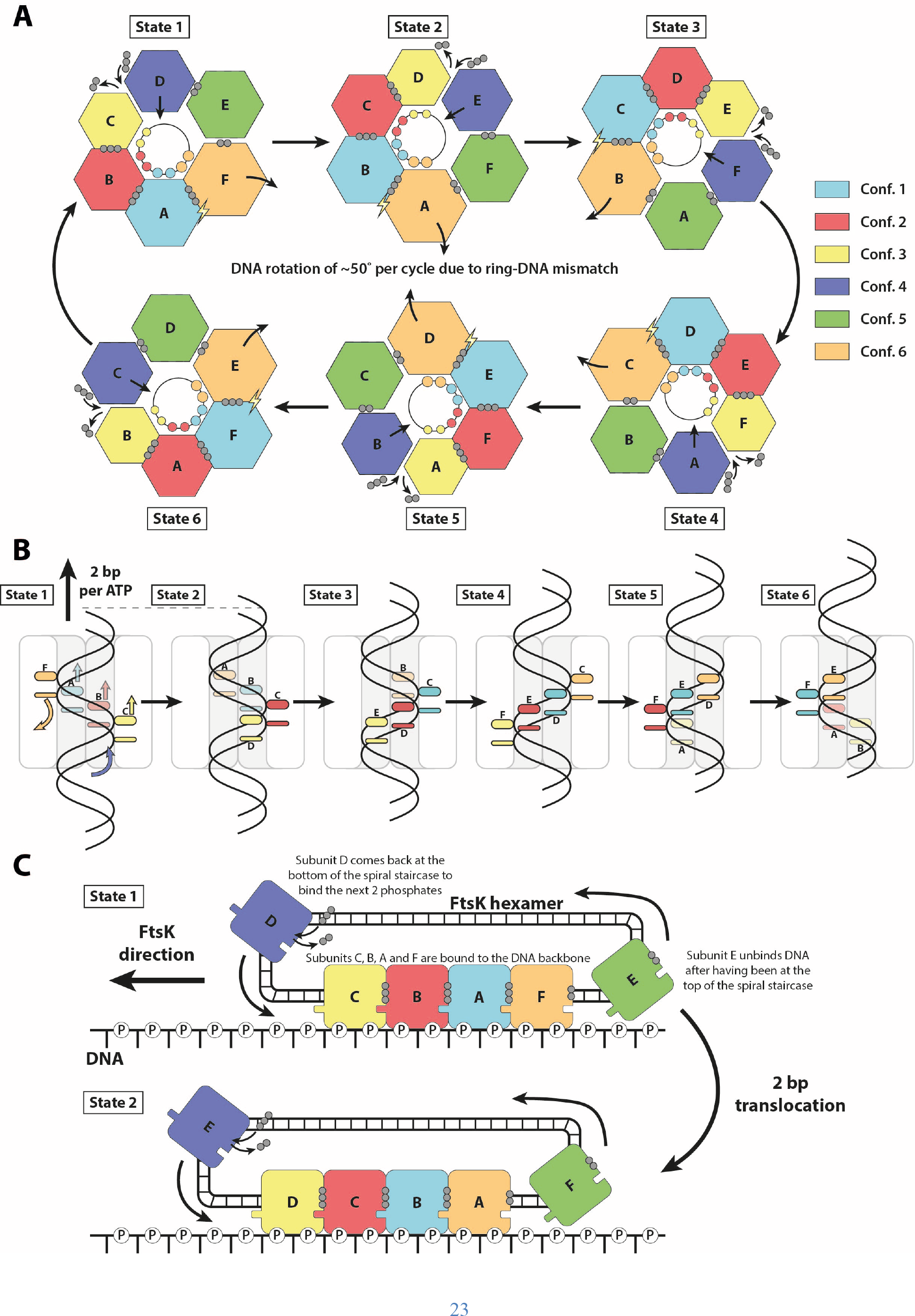
Model for double-stranded DNA translocation. **(A)** The six conformations rotate around the ring. As each subunit advances 1/6th further along the ATPase reaction cycle, it adopts the conformation of the subunit next to it. The conformations, but not the subunits, rotate clockwise around the ring (α-ring view). Because the DNA is helical, rotational adaption of the six conformations leads to 2 bp translocations at each step. Since DNA has 10.5 bp per turn, there is a rotation of FtsK against the DNA of 51.4° per cycle of 6 hydrolysed ATP and 12 bp translocated. Spheres: phosphates; lightning bolt: hydrolysis-competent state. **(B)** DNA translocation through the FtsK_αβ_ pore. In each state, four subunits bind dsDNA through two loops, organized into two spiral staircases. **(C)** The translocation mechanism understood as circularized filament treadmilling. As in cytomotive filaments, the ‘spiral staircase’ along the DNA backbone is extended at one end (conformation 3, yellow, binding DNA next), and shortened at the other end (conformation 6, orange, detaching next). Non-DNA binding conformations 4 & 5 circularize the filament into a ring since they close the conformational wave, resetting conformation 6 into 3.

In each step, all subunits change conformation in a concerted manner, acquiring the next conformation in the cycle. As a result, the subunit at the top of the spiral staircase (subunit F) disengages from dsDNA, switching from conformation 6 to 5. At the bottom of the staircase, another subunit contacts the dsDNA phosphodiester backbone and binds ATP, shifting from conformation 4 to 3 (subunit D in the structure). Every catalytic cycle adds one ATP-bound subunit at the bottom end of the spiral staircase, while removing one ADP-bound at the top, similar to treadmilling of cytomotive protein filaments, but in a closed circular arrangement (Fig. 4C). Using the treadmilling analogy, conformations 4 and 5, resetting the β subdomains from the top to the bottom for renewed binding, represent the recycling of subunits from one end to the other.

The arginine finger, as in other ring-shaped ATPases of similar types (*24*) ensures coordination between the different states. The concerted conformational changes within the spiral staircase all participate in DNA translocation, with each of the four subunits remaining associated to the same two phosphates during the process (Fig. 4B). In our model, two bp of the DNA are translocated per catalytic step and 12 bp by a full cycle around the ring (Fig. 4B), as postulated before (*13*, *22*, *25*). This must be accompanied by a 51.4° anticlockwise rotation of dsDNA per cycle and 12 bp (∼ 4° / bp), and hence, supercoiling (or protein rotation) to compensate for the mismatch with the roughly 10.5 bp per canonical DNA turn.

The α-ring, although probably not actively involved in translocation, may be essential for the high processivity observed by FtsK. We propose that its almost symmetrical arrangement throughout the ring and reaction cycle ensures that the ring does not open during translocation. In addition, it seems to guide the positioning of dsDNA through the pore for optimal interactions with the spiral staircases.

Our model for dsDNA translocation by FtsK_αβ_ recapitulates important features of previous mechanistic models for ring-shaped helicases that act as single-stranded DNA motors, and in particular from the superfamily 5 such as the Rho helicase (*17*, *23*). However, the double-stranded nature of the much more rigid dsDNA substrate of FtsK requires much larger movements of the ATPase subdomains and a second spiral staircase. And the great speed of FtsK seems to require an additional processivity device, the non-deformable α ring. Further studies on other members of the FtsK/HerA family of proteins (*26*) will be essential to extend our insights to dsDNA translocases involved in sporulation, conjugation, dsDNA break repair and viral DNA packaging, and to determine which features make FtsK such an outstandingly fast double-stranded DNA motor.

## Acknowledgments

We thank J. Grimmett and T. Darling, S. Chen, G. Cannone, A. Yeates and G. Sharov for help with computing and EM. We thank J. Wagstaff for stimulating discussions and critical reading of the manuscript. We acknowledge access at the UK National Electron Bioimaging Centre (eBIC), EM17434-66.

## Author contributions

NLJ purified the protein, collected and processed cryo-EM data, and built and refined the structures. TJR acquired the NMR data. TJR and NLJ analysed the NMR data. NLJ and JL wrote the manuscript.

## Data and materials availability

Maps and atomic models of FtsK_αβ_-DNA complexes were deposited in the EM Data Bank (EMDB) and Protein Data Bank (PDB) with accession numbers EMD-10399 and PDB 678B for the ATPγS sample, translocating; EMD-10400 and PDB 678G for the ATPγS sample, stalled; EMD-10402 and PDB 678O for the ATPγS sample, stalled state at DNA end; EMD-10403 (nucleotide-free); EMD-10404 (ADP) and EMD-10405 (AMPPNP). All other data is available in the main text or the supplementary materials.

