## Supplementary Materials for "FtsK in motion reveals its mechanism for double-stranded DNA translocation"

**This PDF file includes:**

Supplementary Materials and Methods
Supplementary Figures S1 to S9
Supplementary Tables S1 and S2
Captions for Supplementary Movies S1 and S2
Supplementary References

**Other Supplementary Materials for this manuscript include the following:**

Movies S1 and S2

### Materials and Methods

#### Cloning and expression

The gene coding for FtsK from *Pseudomonas aeruginosa* strain ATCC 15692 (residues 247 to 728, UniProt entry Q9I0M3) was amplified and cloned into vector pHis17 ahead of a C-terminal KLHHHHHH tag by Gibson assembly (*1*). *Escherichia coli* C41(DE3) cells were transformed by electroporation with the resulting plasmid and plated on a TYE agar plate supplemented with 100 $\mu g.mL^{-1}$ ampicillin. Then, a 50 mL 2xTY pre-culture was grown overnight at 37 ˚C from a single colony in presence of 100 $\mu g.mL^{-1}$ ampicillin. After centrifugation (10 min, 3,220 g, 4˚C), the cells were used to inoculate 6 L of 2xTY medium supplemented with 100 $\mu g.mL^{-1}$ ampicillin. The culture was grown at 37 ˚C until an $OD_{600}$ of approximately 0.6 was reached. Protein expression was induced with 1 mM isopropyl β-d-1-thiogalactopyranoside (IPTG; Anatrace) at 25 ˚C for 6 hours. The cells were finally harvested by centrifugation (20 min, 5,300 g, 4 ˚C) and stored at -20 ˚C.

#### Purification

The cell pellet was resuspended in 150 mL buffer (50 mM Tris, 150 mM NaCl, pH 7.0), complemented with DNase I (Sigma), RNase A (Sigma) and EDTA-free protease inhibitor tablets (Roche). Cells were lysed with a cell disruptor (Constant Systems) at 25 KPSI and the lysate was centrifuged 45 min at 142,000 g in a Type 45 TI rotor (Beckman), at 4 ˚C. 50 mM imidazole was added to the supernatant before loading onto a 5 mL HisTrap HP column (GE Healthcare). The column was washed with buffer A1 (50 mM Tris, 150 mM NaCl, 50 mM imidazole, pH 7.0) and the protein was eluted with step-wise increments of buffer A2 (50 mM Tris, 150 mM NaCl, 1 M imidazole, pH 7.0). Fractions containing the protein were pooled and loaded onto a 5 mL HiTrap Heparin HP column (GE Healthcare). The column was washed with buffer B1 (50 mM Tris, 150 mM NaCl, pH 7.0) before eluting the protein with step-wise increments of buffer B2 (50 mM Tris, 1 M NaCl, pH 7.0). Fractions were pooled, concentrated to 2 mL in a Vivaspin 20 concentrator (PES membrane, 50 kDa molecular weight cut-off, Sartorius). Next, the concentrate was loaded onto a Sephacryl 16/60 S-300 HR column (GE Healthcare) and eluted in buffer C (25 mM Tris, pH 7.5). The protein was concentrated with a Vivaspin 2 concentrator (PES membrane, 50 kDa molecular weight cut-off, Sartorius) before being flash-frozen in liquid nitrogen for storage at -80 ˚C.

#### NMR spectroscopy

1D $^{31}P$ spectra were recorded at 202.4 MHz, 298 K on an Avance II-500 spectrometer (Bruker) equipped with a broadband cryoprobe. Data acquisition time was 3.3 minutes in each spectrum, with 2.9 s inter-scan delay and 0.15 Hz/point digital resolution in the processed data. For each reaction, samples of 1.5 µM $FtsK_{\alpha\beta}$, 180 nM DNA, in 25 mM Tris, 5% (v/v) $D_2O$, pH 7.5 were prepared. Reactions were started with the addition of 1 mM nucleotide and 2 mM $MgCl_2$ (or 2 mM nucleotide and 4 mM $MgCl_2$ for time series). Under these conditions, ATP was fully hydrolysed by $FtsK_{\alpha\beta}$ during the seven minutes dead time between sample insertion and the end of data collection of the first spectrum. Normalised peak integrals for the α-phosphorus measured for the time series spectra were used to estimate the percentage of each nucleotide in the sample over time.

### Cryo-electron microscopy

Mixtures of 0.7 $mg.mL^{-1}$ (13 μM) $FtsK_{\alpha\beta}$, 1.5 μM 45 bp double-stranded (ds) DNA (sequence CGCAGGAAAAATAGCGATTTGAAGGATTCGACCACCGCGAGCCAT) in 25 mM Tris pH 7.5 were prepared on ice. After 2 min incubation with final concentrations of 2 mM ATPγS and 4 mM $MgCl_2$, 3 μL of the samples were added to glow-discharged R1.2/1.3 AU 300 mesh Quantifoil grids. The grids were blotted for 2 sec at force -5 and flash-frozen into liquid ethane using a Thermo Fisher Vitrobot Mark IV. Nucleotide-free, ADP and AMPPNP samples were prepared similarly with incubation times up to 15 min. The $FtsK_{\alpha\beta}$-dsDNA complex in presence of ATPγS was imaged on a Titan Krios electron microscope from the Electron Bio-Imaging Centre (eBIC) at Diamond Light Source. 3,300 micrographs were collected using EPU software on a K2 camera in counting mode at a nominal magnification of 130,000 x, with a resulting pixel size of 1.048 Å. Each movie was recorded for 12 sec, with a total dose of ~ 43 $e^-.Å^{-1}$ over 48 fractions, within a nominal defocus range of -3.5 to -2.0 μm. The other $FtsK_{\alpha\beta}$-dsDNA complex samples (nucleotide-free, ADP and AMP-PNP) were imaged similarly on an in-house Titan Krios microscope. Micrographs were collected on a K2 camera in counting mode, at pixel sizes ranging from 1.08 to 1.145 Å, and total doses from ~ 39 to ~ 49 $e^-.Å^{-1}$.

### Cryo-EM data processing

All data were processed in Relion 3.0 (*2*). For the $FtsK_{\alpha\beta}$-dsDNA-ATPγS sample, dose-weighted micrographs were motion corrected with MotionCor2 (*3*) and their contrast transfer function (CTF) parameters were estimated with CTFFIND 4.1 (*4*). Using an estimated resolution cut-off of 6 Å, 49 images were discarded from the next steps. 414 particles were manually picked and 2D classified. Using these 2D classes as references, a total of 1,510,109 particles were picked automatically and extracted. After 2D classification, 1,094,683 particles were selected and sorted into four classes through 3D classification using as a reference a 40 Å low-passed filtered map of the $FtsK_{\alpha\beta}$-ring (PDB 2IUU, no DNA) and a 180 Å circular mask. While one class was discarded for its poor quality, the three others were separately reconstructed (with a mask including just the FtsK ring and the DNA) and each class was used for a round of further 3D classification without alignment. The resulting classes were again reconstructed, and those with the highest resolutions and best DNA density were further improved using per particle CTF correction, Bayesian polishing and postprocessing. This resulted in three maps, one in a highly asymmetric state (later determined to be translocating) at 3.6 Å resolution (map "III.E", see Fig. S3), and two much more symmetrical states (later determined to be stalled), one of which with dsDNA visible all along the pore (4.34 Å resolution, "III.B") while in the other one dsDNA is found only in the upper half of the pore, towards the pore's exit (3.99 Å resolution, "II.A"). During processing, many other classes appeared that represented only small variations of each of these three, with often less defined DNA or lower resolution for some subunits. These were not used for final refinements as their inclusion decreased quality of the generated maps.

$FtsK_{\alpha\beta}$-dsDNA samples in the nucleotide-free, ADP- or AMPPNP-states were processed similarly in the early stages with 393,540, 240,870, and 265,603 particles selected, respectively, after 2D classification. However, after 3D classification EM maps with a well-resolved symmetrical ring but a featureless dsDNA were obtained for each dataset. Given the small size of the DNA compared to the protein, and the absence of any visible conformational variation among the subunits forming the ring, it is likely that the particles' relative orientation around the (pseudo) six-fold axis of the protein was wrongly estimated. Further processing steps involved $FtsK_{\alpha\beta}$ signal subtraction and 3D classification on the DNA itself (using a 30 Å low-passed filtered map generated from a dsDNA model). Classes with visible DNA grooves were selected (56,904, 29,563, and 28,958 particles for $FtsK_{\alpha\beta}$-dsDNA respectively without any nucleotide, with ADP or with AMP-PNP), and were refined and postprocessed based on the full particles signal, to reconstruct the complexes with only partial recovery of the dsDNA's detailed features.

**Model building and refinement**

The crystal structure of $FtsK_{\alpha\beta}$ (PDB code 2IUU) was docked inside map III.E using UCSF Chimera (*5*) to serve as a starting model for atomic model building of the translocating complex. For the dsDNA, a 20 bp B-form dsDNA model was generated in Coot (*6*) and fitted manually inside the map. Subsequently, the model underwent several rounds of manual rebuilding in Coot and real-space refinement in Phenix (*7*). The model's stereochemistry and fit was validated with Molprobity and EMRinger (*8*, *9*). Similar procedures were performed for $FtsK_{\alpha\beta}$-dsDNA stalled states associated with maps II.A and III.B.

**Model and morph of dsDNA translocating through $FtsK_{\alpha\beta}$**

The six conformations around the $FtsK_{\alpha\beta}$ ring each represent a snapshot of $1/6^{th}$ of the complete hydrolysis cycle of each subunit. They hence represent a wave-like continuous conformational change around the ring. By changing each subunit's conformation into the conformation of the next conformation in the reaction cycle (the subunit next to it in the ring), simultaneously for all subunits, an atomic morph of a complete translocation cycle can be generated through interpolation between each snapshot (six ATP hydrolysed, 12 bp translocated). Briefly, each protein-DNA state was generated by incremental 60° (clockwise from top) rotation of the entire $FtsK_{\alpha\beta}$-dsDNA complex. Subunits were renamed afterwards so that every protomer keeps the same chain ID for a given position in the ring. DNA translocation was performed by extending the dsDNA by 2 bp at one end (exit side), shortening it by the same length on the other side at each step, and renumbering the bases so that each strand's 5' end always starts with base number 1. Intermediate states for Movie S2, between the six key states generated this way were derived by morphing in PyMOL. Together, these steps allowed visualizing the slight rotation of dsDNA as well as its continuous slight deformation as it traverses the channel (see top view in Movie S2).

### Supplementary Figures

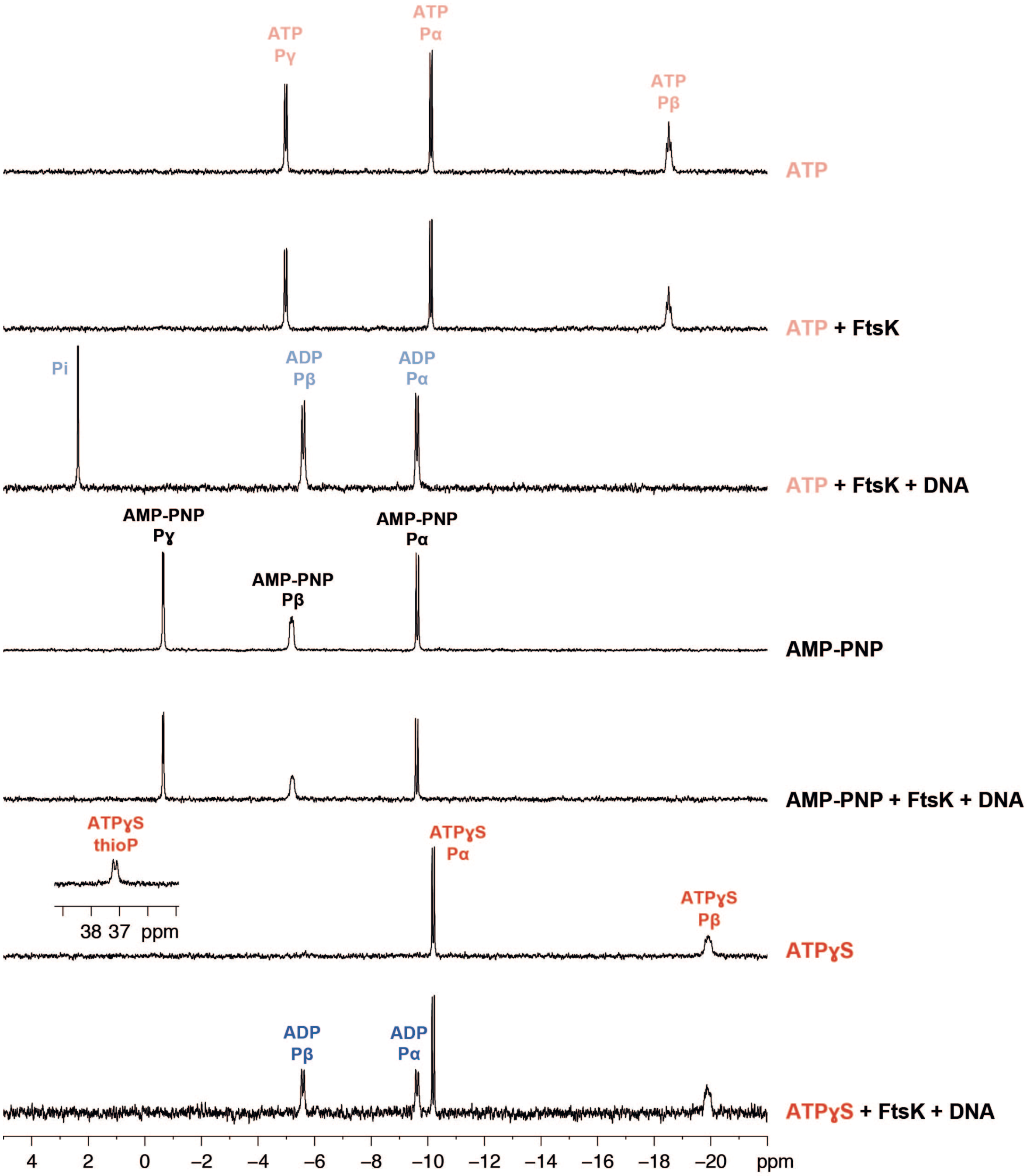

**Figure S1. Nucleotide hydrolysis activity of FtsK$_{\alpha\beta}$ measured by NMR.** 1D $^{31}$P NMR spectra of different mixtures of FtsK$_{\alpha\beta}$, dsDNA and nucleotides. Samples were prepared as detailed in Materials & Methods. Spectra were collected seven minutes after nucleotide addition. ATP (fast) and ATPγS (slow) were hydrolysed by FtsK$_{\alpha\beta}$ only in the presence of DNA, but not AMPPNP.

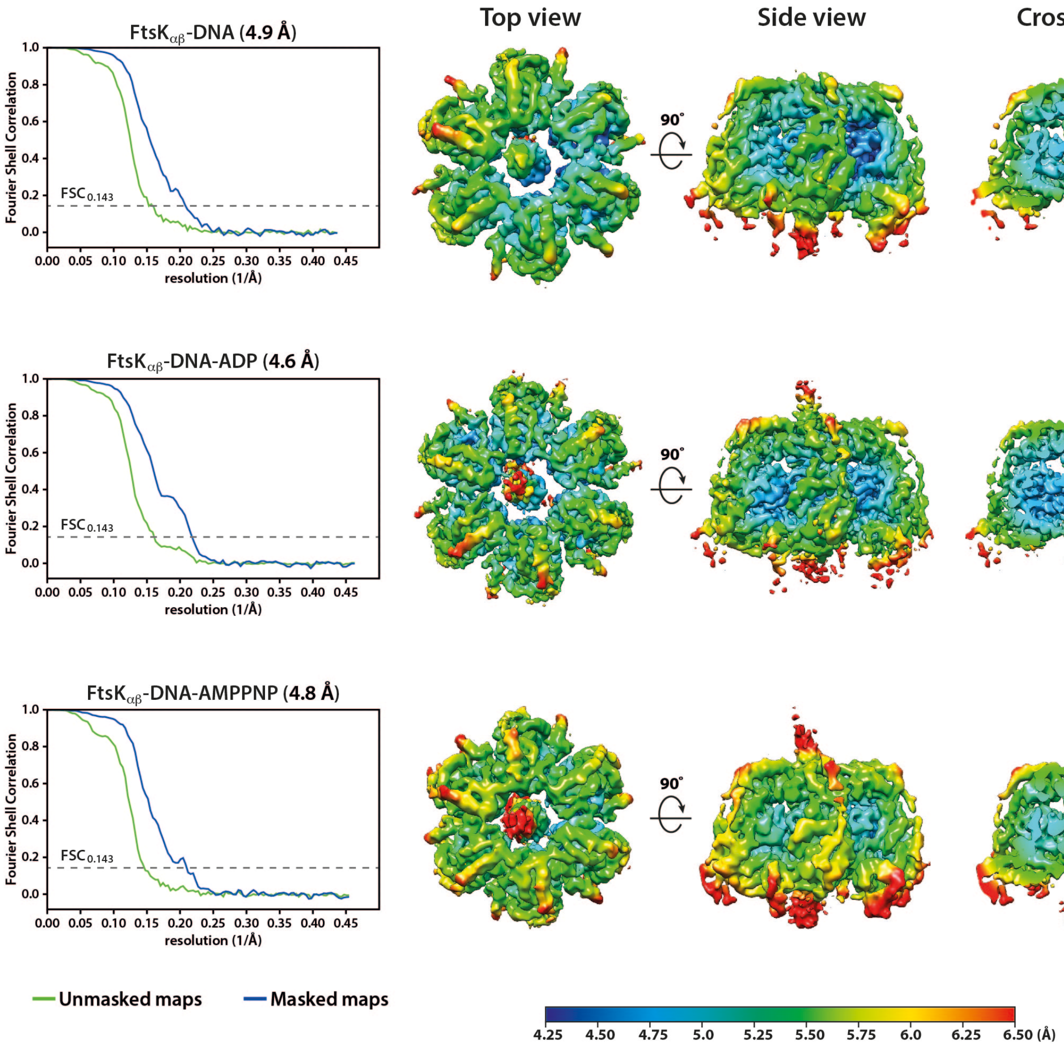

**Figure S2. Resolution estimation of cryo-EM maps for FtsK$_{\alpha\beta}$-DNA complexes in the nucleotide-free, ADP- and AMPPNP- states.** Left panels show the Fourier shell correlation (FSC) curves for masked (blue) and unmasked (green) maps using gold standard half-maps. The $FSC_{0.143}$ threshold for resolution estimation is indicated by a dotted grey line. The panels on the right show top, side and cross-sectioned views of the complexes, coloured by local resolution as determined by Relion 3.0.

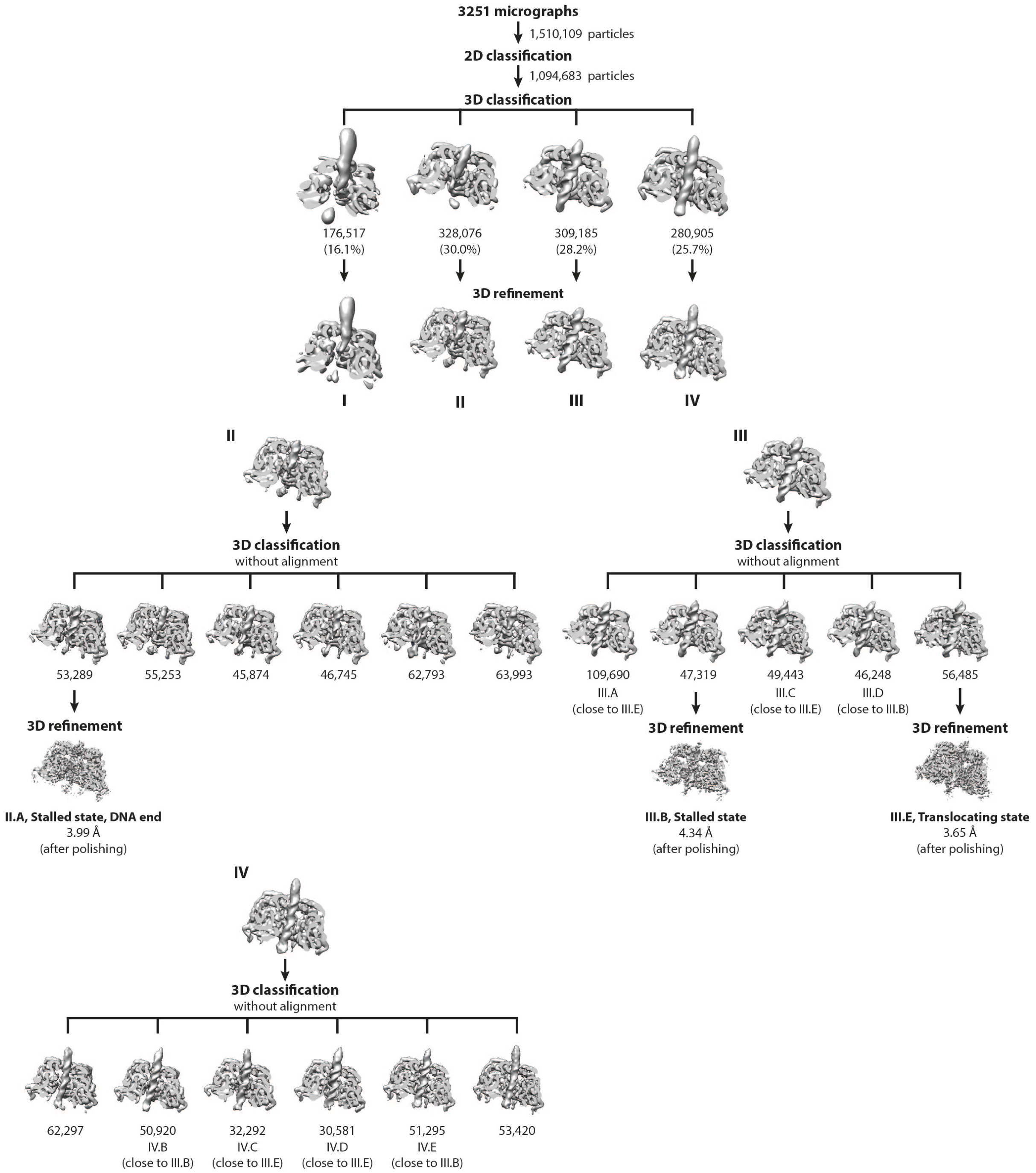

**Figure S3. Data processing workflow of the FtsK$_{\alpha\beta}$-dsDNA + ATPγS dataset.** Processing was performed in Relion 3.0. After 3D classification, good particles were sorted into three main classes (II, III and IV). Further 3D classification identified three classes (II.A, III.B and III.E) that were better resolved and had very well-defined DNA density. The number of particles in each class is indicated underneath.

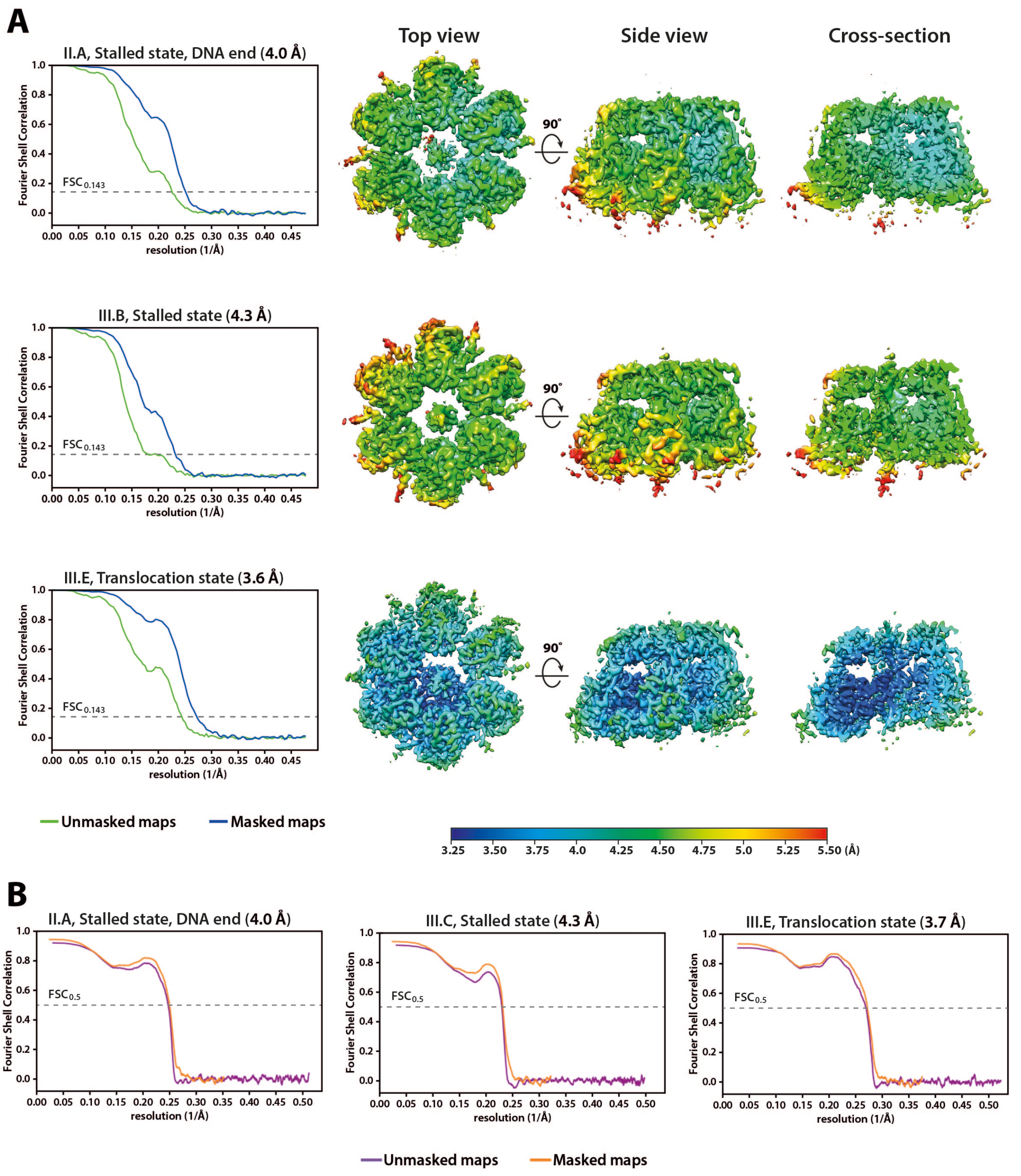

**Figure S4. Resolution estimation of cryo-EM maps and atomic models for the FtsK$_{\alpha\beta}$-dsDNA complex in the ATPγS sample. (A)** Resolution estimation of FtsK$_{\alpha\beta}$-dsDNA complexes. Left panels show the Fourier shell correlation (FSC) curves for masked (blue) and unmasked (green) maps using gold standard half-maps. The FSC$_{0.143}$ threshold for resolution estimation is indicated by a dotted grey line. Panels on the right represent top, side and cross-sectioned views of the complexes, coloured by local resolution as determined by Relion 3.0. (**B**) Map-to-model FSCs.

Curves in orange are for masked maps and purple curves represent unmasked maps. The $FSC_{0.5}$ threshold is represented by a dotted grey line. Resolutions estimated from each FSC curves are indicated above and closely match those estimated for the maps the models are derived from and have been refined against.

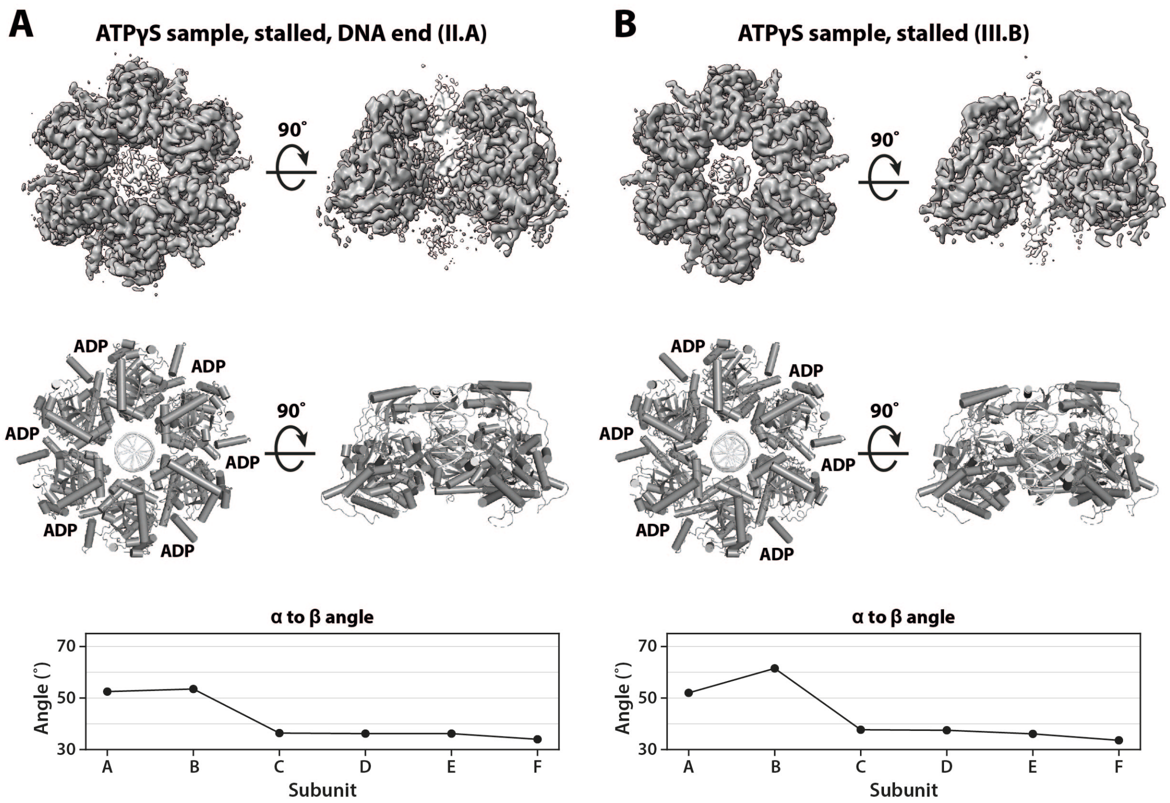

**Figure S5. Further structures of $FtsK_{\alpha\beta}$-dsDNA complexes from the ATPγS sample. (A)** and **(B)** show maps and atomic models of $FtsK_{\alpha\beta}$ with partial (DNA end) or full DNA signals, respectively. The graphs in lower panels illustrate the angles measured between the α and β subdomains across the 6 subunits. Because the structures do not contain nucleotide states of a full hydrolysis cycle around the ring, the angle distribution does not show a wave function (as in Figure 2A, bottom; and hence the ring is much less asymmetric) and the DNA density is truncated and/or less well defined, we propose that these structures represent stalled states of the motor.

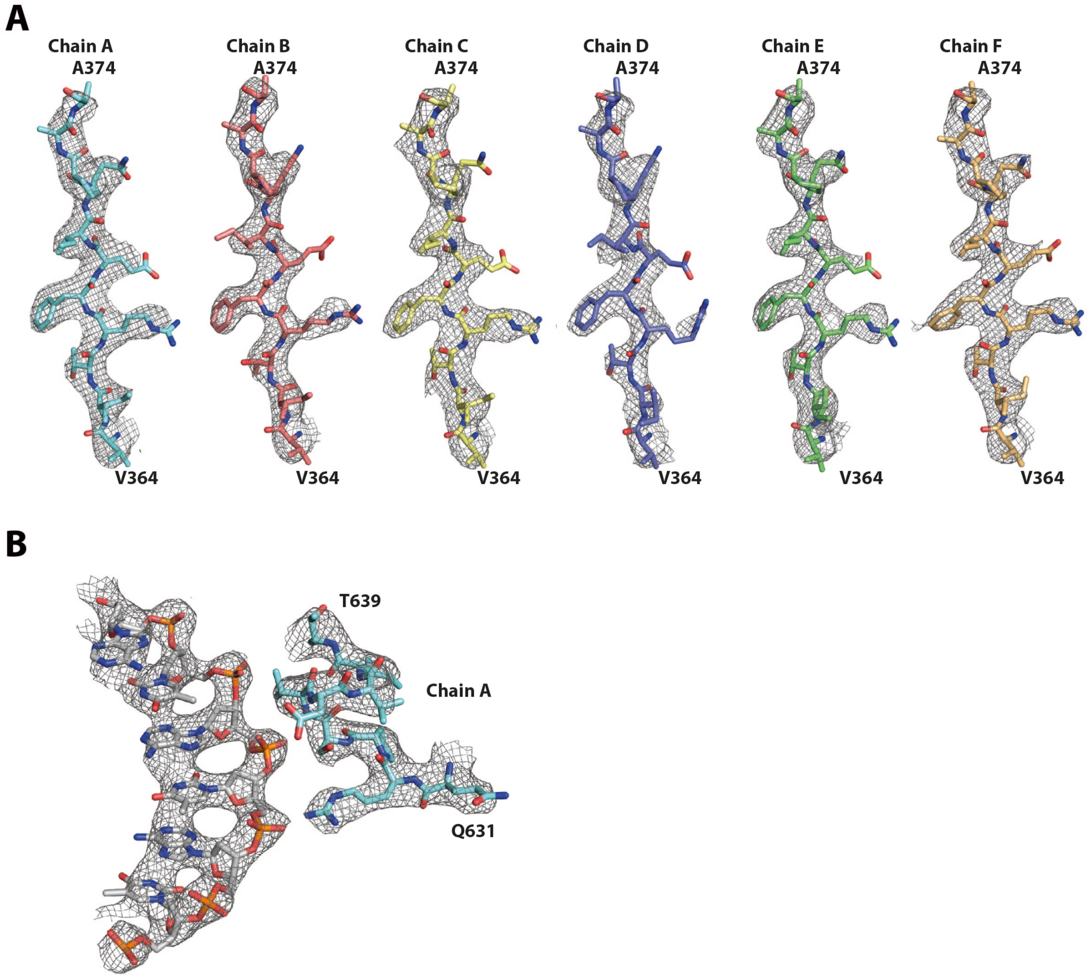

**Figure S6. Representative cryo-EM map details of the translocating FtsK$_{\alpha\beta}$-dsDNA complex in the translocating state. (A)** Residues V364 to A374 of all six subunits and **(B)** the interaction between loop I from chain A and one DNA strand. The refined atomic models for these regions are shown superimposed.

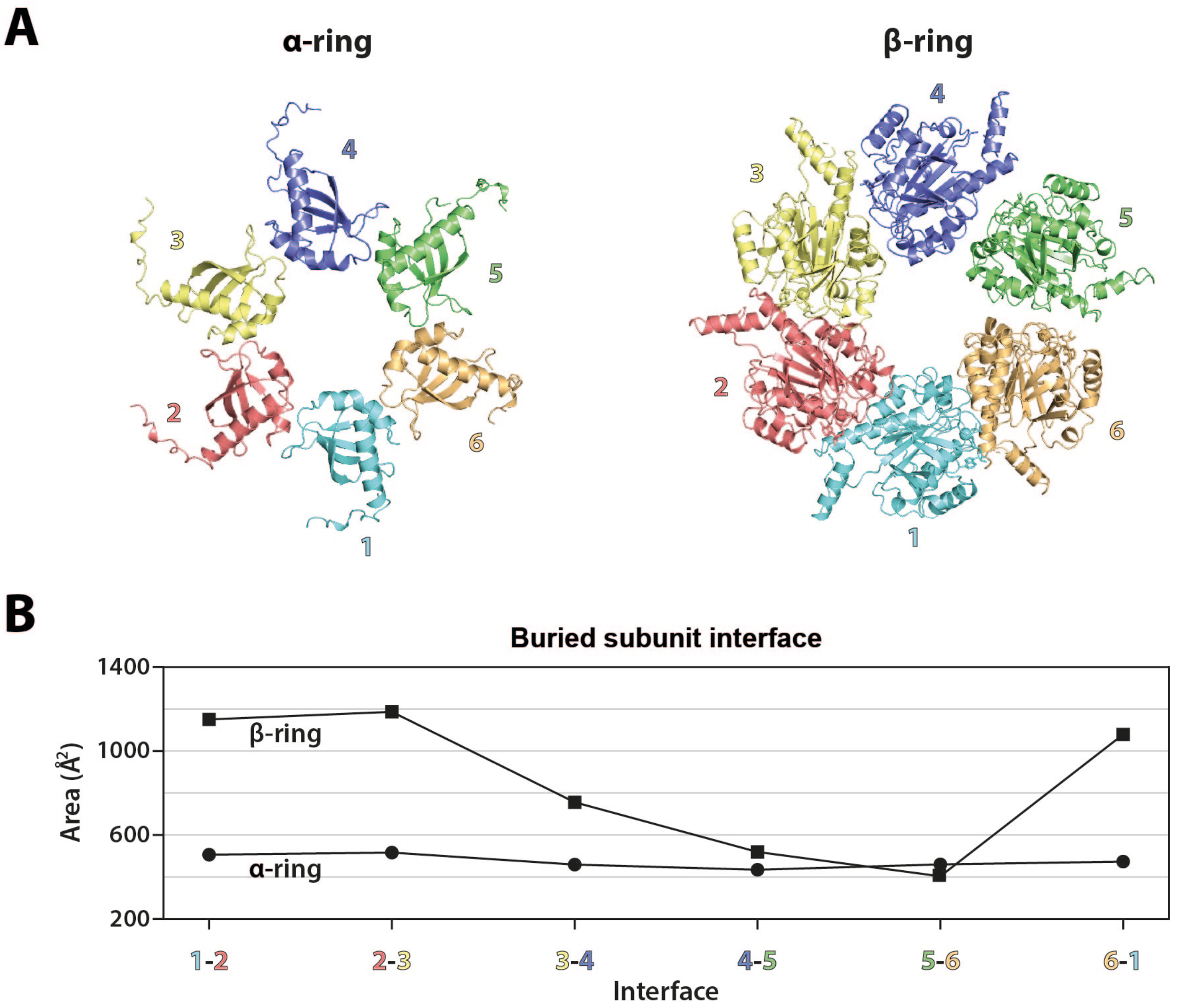

**Figure S7. Molecular packing analysis of the translocating FtsK$_{\alpha\beta}$-dsDNA complex in the ATPγS sample. (A)** Structures of the isolated α (left) and β (right) rings. Subunit conformations are indicated by numbers shown next to each structure. **(B)** Buried subunit surfaces in the interfaces along each ring. Note that the α ring is more-or-less symmetric, whereas the β ring is not.

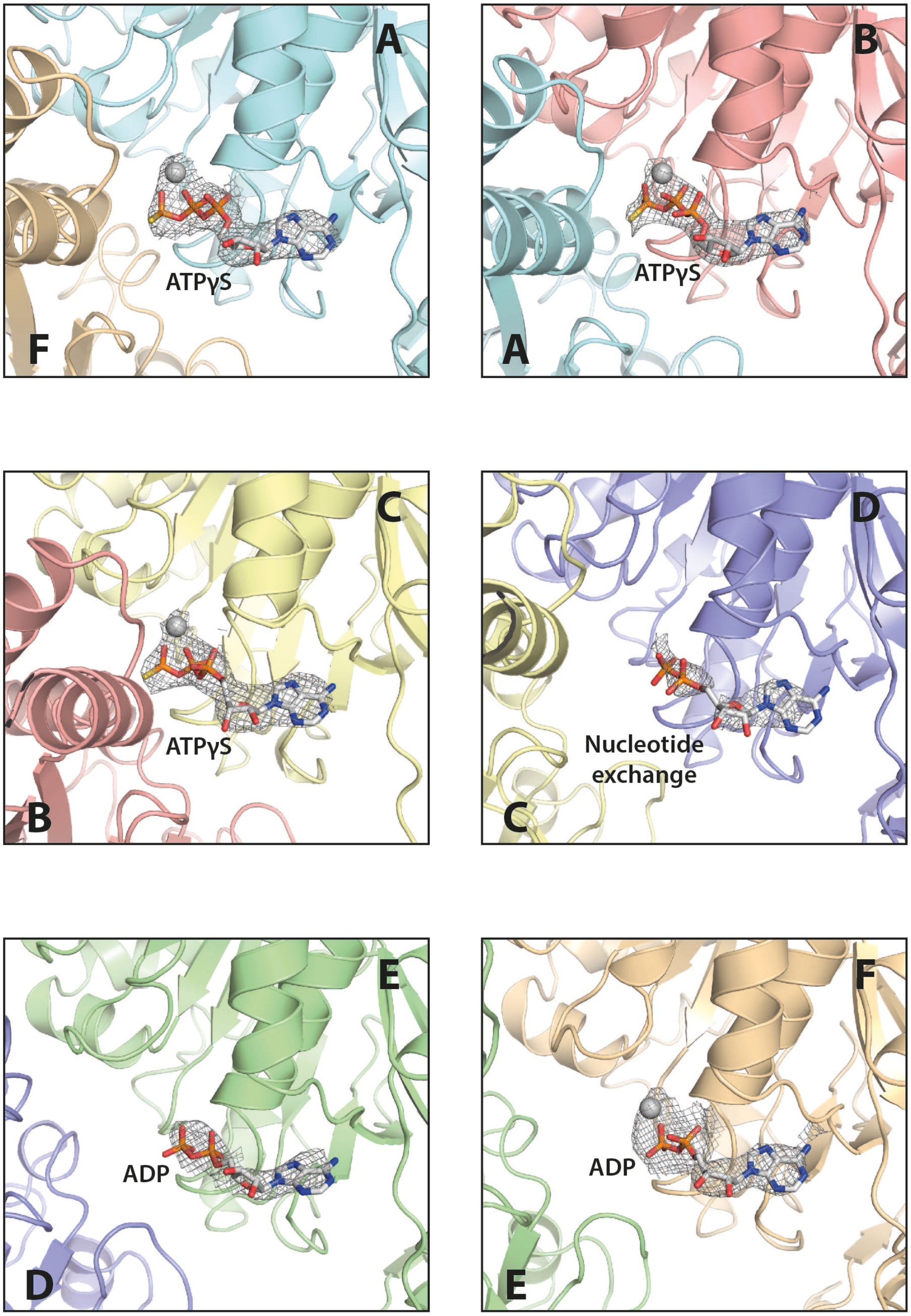

**Figure S8. Cryo-EM map details around the nucleotides in the FtsK$_{\alpha\beta}$-dsDNA complex (translocating state).** Superposition of the nucleotide cryo-EM maps and the atomic model of the assigned nucleotides.

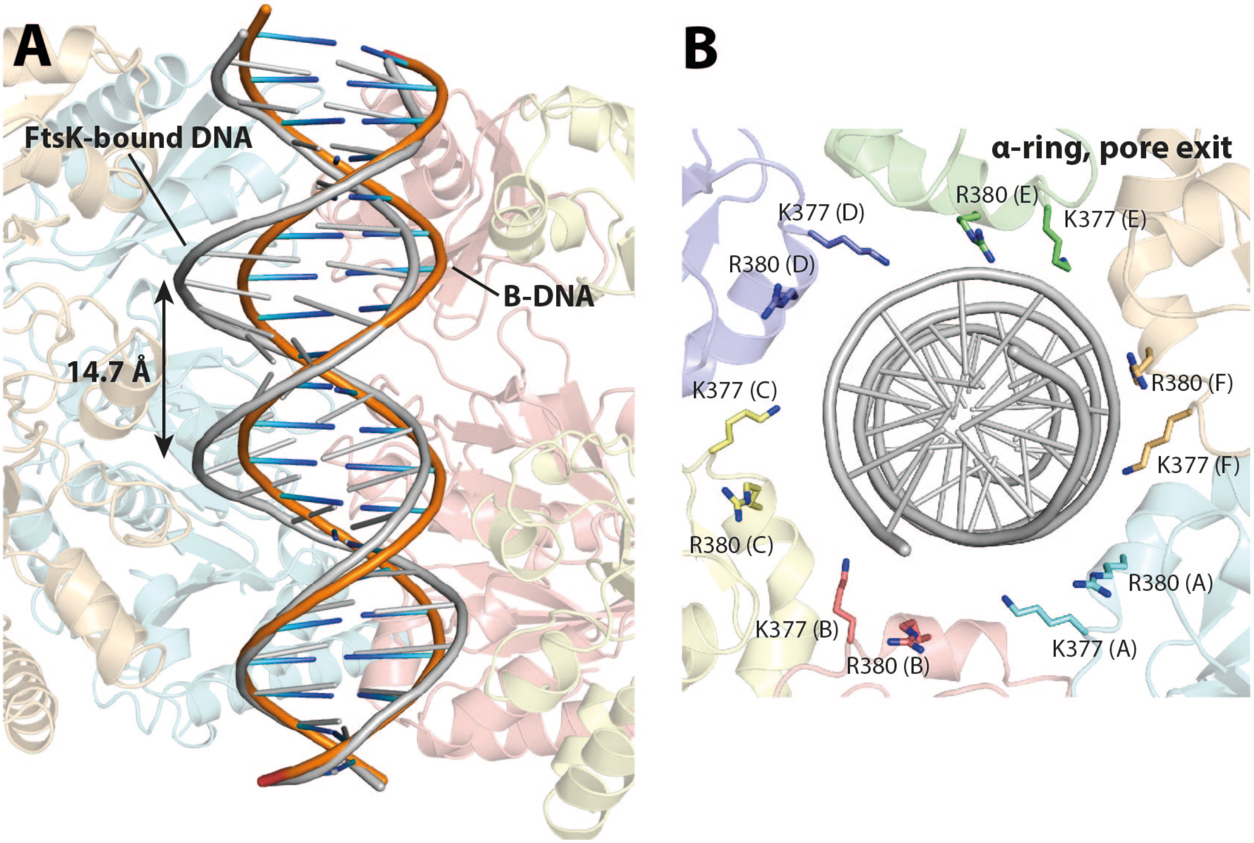

**Figure S9. Atomic details of FtsK$_{\alpha\beta}$-dsDNA interactions in the translocating state.** (A) DNA distortion by FtsK$_{\alpha\beta}$. Interaction with FtsK$_{\alpha\beta}$ widens the DNA's minor groove by up to 2.9 Å (+25%) (grey) compared to canonical B-form DNA (orange). **(B)** Top view of DNA pore formed by the α-ring. Residues K377 and R380 on each subunit create a positively charged ring at the pore's exit, shown here as sticks.

### Supplementary Tables

#### Table S1. Cryo-EM data collection and processing of nucleotide-free, ADP- and AMPPNP-bound $FtsK_{\alpha\beta}$-dsDNA complexes.

| | **$FtsK_{\alpha\beta}$-DNA (EMD 10403)** | **$FtsK_{\alpha\beta}$-DNA-ADP (EMD 10404)** | **$FtsK_{\alpha\beta}$-DNA-AMPPNP (EMD 10405)** |
|---|---|---|---|
| **Data collection** | | | |
| Microscope | Titan Krios | Titan Krios | Titan Krios |
| Voltage (kV) | 300 | 300 | 300 |
| Detector | K2 Summit | K2 Summit | K2 Summit |
| Nominal magnification | 105,000 | 130,000 | 105,000 |
| Pixel size (Å) | 1.145 | 1.08 | 1.10 |
| Total electron fluence (e-/Å$^2$) | 43.8 | 39.4 | 48.6 |
| Defocus range (μm) | -3.5 to -2 | -3.5 to -2 | -3.5 to -2 |
| **Data processing** | | | |
| Micrographs | 742 | 771 | 830 |
| Extracted particles | 397,748 | 514755 | 311,615 |
| Refined particles | 393,540 | 240,870 | 265,603 |
| Final particles | 56,904 | 29,563 | 28,958 |
| Map resolution (Å) | 4.91 | 4.63 | 4.80 |
| FSC threshold | 0.143 | 0.143 | 0.143 |
| Map resolution range (Å) | 4.35 to 7.0 | 4.55 to 7.0 | 4.6 to 7.1 |

**Table S2. Cryo-EM data collection, processing and refinement statistics of structures from FtsK$_{\alpha\beta}$-dsDNA plus ATPγS.**

| | PaFtsK$_{\alpha\beta}$-DNA-ATPγS Translocation state (EMD 10399, PDB 6T8B) | PaFtsK$_{\alpha\beta}$-DNA-ATPγS Stalled state (EMD 10400, PDB 6T8G) | PaFtsK$_{\alpha\beta}$-DNA-ATPγS Stalled state, DNA end (EMD 10402, PDB 6T8O) |
|---|---|---|---|
| **Data collection** | | | |
| Microscope | Titan Krios | Titan Krios | Titan Krios |
| Voltage (kV) | 300 | 300 | 300 |
| Detector | K2 Summit | K2 Summit | K2 Summit |
| Nominal magnification | 130,000 | 130,000 | 130,000 |
| Pixel size (Å) | 1.048 | 1.048 | 1.048 |
| Total electron fluence (e-/Å$^2$) | 42.95 | 42.95 | 42.95 |
| Defocus range (µm) | -3.5 to -2 | -3.5 to -2 | -3.5 to -2 |
| **Data processing** | | | |
| Micrographs | 3,300 | 3,300 | 3,300 |
| Extracted particles | 1,510,109 | 1,510,109 | 1,510,109 |
| Refined particles | 1,094,683 | 1,094,683 | 1,094,683 |
| Final particles | 56,485 | 47,319 | 53,289 |
| Map resolution (Å) | 3.65 | 4.34 | 3.99 |
| FSC threshold | 0.143 | 0.143 | 0.143 |
| Map resolution range (Å) | 3.4 to 5.0 | 4.1 to 5.8 | 3.8 to 5.4 |
| **Refinement** | | | |
| Initial model used (PDB code) | 2IUU | 2IUU | 2IUU |
| Map sharpening B-factor (Å$^2$) | -90.9 | -147.3 | -120.1 |
| Model composition | | | |
| Non-hydrogen atoms | 19,077 | 18,875 | 18,806 |
| Residues | 2370 (FtsK); 40 (DNA) | 2367 (FtsK); 32 (DNA) | 2374 (FtsK); 26 (DNA) |
| Ligands | 10 (6 nucleotides, 4 ions) | 6 | 6 |
| Map correlation coefficient | 0.82 | 0.83 | 0.82 |
| R.m.s deviations | | | |
| Bond lengths (Å) | 0.004 | 0.008 | 0.006 |
| Bond angles (˚) | 0.720 | 1.138 | 1.242 |
| **Validation** | | | |
| MolProbity score | 2.11 | 1.95 | 1.61 |
| Clashscore | 13.97 | 7.19 | 4.39 |
| Poor rotamers (%) | 0.0 | 0.15 | 0.20 |
| Ramachandran plot | | | |
| Favored (%) | 92.80 | 89.80 | 94.21 |
| Allowed (%) | 7.20 | 10.20 | 5.79 |
| Disallowed (%) | 0.00 | 0.00 | 0.00 |
| EMRinger score | 2.47 | 1.45 | 1.99 |

**Legends for Supplementary Movies**

**Movie S1. Recognition of dsDNA by a double 'spiral staircase' of DNA-interacting loops is facilitated by FtsK$_{\alpha\beta}$'s conformational diversity around the ring.** Each subunit is coloured differently according to their conformation. The translocating III.E structure contains three distinct nucleotide states (ATPγS, ADP, and nucleotide exchange). Basic residues from loops I and II are interacting with the minor groove of the double-stranded DNA (interacting residues coloured in red) and are arranged as two spiral staircases containing four of the six subunits. Disengaged residues (K643 from conformation 1, residues from subunits D and E) are depicted in grey.

**Movie S2. Model for dsDNA translocation by FtsK$_{\alpha\beta}$.** The movie shows the concerted conformational changes during the catalytic cycle to translocate dsDNA. Each subunit goes through all six stages of hydrolysis and their associated conformational states, meaning that the conformations rotate around the ring (depicted by colours). The DNA rotates only slowly against the subunits because of a symmetry mismatch between the 10.5 bp per turn of DNA and the 12 bp translocation step (6 times 2) per full cycle. In the movie, translocation is first shown from the top (looking towards the α-subdomains), where the slow rotation of dsDNA due the DNA-hexamer base pair mismatch can be seen. In the following side view, one and later on five subunit(s) is (are) hidden for clarity. Subunit colours depict conformations throughout and correspond to Figure 4. Atomic motions are interpolations (morphs) between the six conformations observed in the translocating state III.E.
